# ESAT-6 protein of *Mycobacterium tuberculosis* inhibits differentiation of human monocytes to dendritic cells

**DOI:** 10.1101/2025.08.06.668872

**Authors:** Akshay Girish Manikoth, Rahila Qureshi, Sangita Mukhopadhyay

**Author notes:** Corresponding author **Address correspondence to** Sangita Mukhopadhyay, Laboratory of Molecular Cell Biology, BRIC-Centre for DNA Fingerprinting and Diagnostics, Inner Ring Road, Uppal, Hyderabad– 500039, India. Equal contribution (Akshay Girish Manikoth, Rahila Qureshi).

## Abstract

*Mycobacterium tuberculosis* (Mtb) employs multiple strategies to evade host immunity, including disruption of antigen presentation. Dendritic cells (DCs) are crucial for effective antigen presentation and T-cell activation. In this study, we show that mycobacterial protein, ESAT-6, impairs monocyte to DC differentiation with reduced expression of the DC marker, CD209. ESAT-6 treatment elevated IL-6 and IL-10 levels, but blocking of these cytokines failed to restore DC differentiation. Mechanistically, ESAT-6 suppressed phosphorylation of p65, establishing that ESAT-6 impairs DC differentiation by inhibiting NF-κB activation and this function is dependent on the last six amino acids of its C-terminal domain. This mechanism may represent a novel immune evasion strategy employed by Mtb to subvert host adaptive immune responses during infection.

## Introduction

Tuberculosis (TB), caused by *Mycobacterium tuberculosis* (Mtb), remains one of the leading causes of death worldwide, holding the status of the top infectious killer globally [1]. Mtb has evolved numerous strategies to evade host immune responses to favor its persistence and replication [2]. The bacilli are shown to evade antigen presentation pathways [2] and inhibit differentiation of dendritic cells (DCs) to dampen host’s adaptive immune responses [3]. DCs are professional antigen-presenting cells (APCs) that are responsible for capturing and processing antigens for presenting them to T-cells [4]. This results in activation of CD4^+^ and CD8^+^ T-lymphocytes and B-cells, culminating in a robust adaptive immune response that activate immune defense against Mtb [4–6].

Although DCs originate primarily from bone marrow precursors, they can also be differentiated from peripheral blood monocytes upon exposure to cytokines such as granulocyte-macrophage colony-stimulating factor (GM-CSF) and interleukin-4 (IL-4) [7]. Monocyte-derived dendritic cells (MoDCs) are shown to activate the protective Th1-type responses at the site of infection [8]. MoDCs exhibit functional capabilities comparable to those of classical dendritic cells, including the cross-presentation of protein antigens. Circulating monocytes serve as a key reservoir that can rapidly generate DC-SIGN dendritic cells in response to microbial stimuli, enabling effective antigen presentation and T-cell activation [9]. MoDCs play essential roles in both innate and adaptive immunity by initiating CD4 and CD8 T-cell responses, supporting immunoglobulin production by B-cells, and contributing to early immune defense [10]. Monocytes also serve as precursors for DCs residing in peripheral tissues specialized for antigen capture and presentation [8].

While it is established that Mtb impairs differentiation of human monocytes into dendritic cells [3], the specific bacterial factors mediating this effect remain poorly understood. One of the virulence factors of Mtb that is known to subvert adaptive immune responses of host is the early secreted antigenic target of 6 kDa (ESAT-6), encoded by the region of difference 1 (RD1) of the Mtb genome [11]. ESAT-6 is secreted by *M. tuberculosis*, and its deletion results in significant attenuation of mycobacterial virulence [12]. ESAT-6 has been implicated in a range of host-modulatory functions, including inhibition of IL-12 production [13], M2 polarization of macrophages [14], interference with phagosome maturation [15], disruption of autophagy (16), induction of apoptosis in infected macrophages [17], and inhibition of MHC class I antigen presentation [18]. Since ESAT-6 plays a crucial role in Mtb virulence and is shown to modulate host adaptive immune responses [11], it is possible that Mtb uses ESAT-6 protein to inhibit differentiation and function of MoDC to escape sterilizing immunity. In this study, we demonstrate that ESAT-6 significantly impairs MoDC differentiation by targeting NF-κB signaling. Our findings reveal a novel role for ESAT-6 in subverting host adaptive immunity. These insights are likely to deepen our understanding of Mtb immune evasion and may provide new targets for host-directed TB therapies.

## Materials and Methods

### Expression and purification of recombinant proteins

The ESAT-6 and C-terminal deleted ESAT-6 (ΔESAT-6) were purified as described earlier [18]. Briefly, purification was carried out under denaturing condition using urea employing Ni-NTA affinity chromatography and refolding by gradual removal of urea. Protein purity was confirmed by Tris-Tricine SDS-PAGE, and LPS contamination was removed using polymyxin B-agarose beads. The purified proteins were dialyzed, quantified, and filter-sterilized. These preparations were used in subsequent experiments.

### Cell culture for differentiation of monocytes to dendritic cells

Human peripheral blood mononuclear cells (PBMCs) were purchased from HiMedia, India and monocytes were isolated *via* CD14^+^ selection using magnetic activated cell sorting (MACS) system (Stem Cell Technologies, Canada). Monocytes were cultured in complete RPMI-1640 with growth factors, human recombinant GM-CSF (100 ng/ml) and human recombinant IL-4 (50 ng/ml) (PeproTech, ThermoFisher Scientific, USA) for 7 days, with or without ESAT-6 or ΔESAT-6. In some cultures, we added either of the pharmacological inhibitor *viz.* SB203580, S3I-201, BAY 11-7082, wortmannin, or neutralizing antibodies for IL-6 and IL-10 (Invitrogen, ThermoFisher Scientific, USA). Supernatants were collected on days 4 and 7 for ELISA, and cells were harvested on day 7 for flow cytometry.

### Flow cytometry analysis

For flow cytometry analysis, cells were washed and blocked with bovine serum albumin (BSA) and then incubated with either CD209-FITC antibody (BD Pharmingen™) in FACS staining buffer (1X PBS containing 1% BSA and 0.1% sodium azide) for 60 minutes on ice. Cells were washed thrice with FACS staining buffer and cell-bound fluorescence was measured on FACSAria III flow cytometer (Beckton Dickinson, USA). The FACS data were analyzed using FlowJo data analysis software (TreeStar, Oregon, USA).

### Western blot analysis

Western blotting was performed as described earlier [19]. Briefly, cells were lysed in RIPA buffer with phosphatase and protease inhibitors, and debris was cleared by centrifugation. Samples were resolved by SDS-PAGE and transferred to PVDF membranes. Membranes were probed with antibodies against total or phospho-p65 NF-κB or β-actin (Cell Signaling Technology, USA), followed by HRP-labeled anti-Rabbit IgG. Bands were visualized using chemiluminescence.

### Enzyme-linked immunosorbent assay (ELISA)

Levels of IL-6 and IL-10 cytokines in the culture supernatants were measured by ELISA using a commercially available ELISA kit (Invitrogen, ThermoFisher Scientific, USA) following the manufacturer’s protocol.

### Statistical Analysis

GraphPad 9.0 Prism software was used for determining the significance of difference between the groups. Statistical significance was determined using one-way analysis of variance (ANOVA), followed by Tukey’s multiple comparison test. p <0.05 was considered to be significant.

## Results

### ESAT-6 inhibits differentiation of human monocytes into dendritic cells

To examine the effect of ESAT-6 on differentiation of monocyte-derived dendritic cells (MoDCs), human monocytes were treated with recombinantly purified ESAT-6 protein for 30 minutes prior to the addition of GM-CSF and IL-4 (GM-CSF+IL-4). It is known that GM-CSF+IL-4 is the commonly used cytokine combination that stimulates *in vitro* differentiation of monocytes into DCs [20]. Cultures were maintained for 7 days at 37°C, and then the morphological and phenotypical status of cells was assessed. Microscopic analysis revealed that the GM-CSF+IL-4 culture had cells with characteristic dendritic morphology, including elongated shapes, dendritic projections, and the formation of loosely associated cellular clusters, indicating that GM-CSF+IL-4 caused differentiation towards a DC-phenotype, which was expected [20] (Fig. 1A). In contrast, ESAT-6–treated cells failed to acquire dendritic morphology. These cells remained rounded, lacked dendritic extensions, and did not form the typical clusters observed in differentiating DC cultures (Fig. 1A). Flow cytometric analysis was then performed to assess the expression of CD209 (DC-SIGN), a surface marker associated with MoDCs [9]. Cells treated with ESAT-6 displayed a significant reduction in CD209 expression compared to those treated with GM-CSF+IL-4 alone (Fig. 1B). Notably, CD14 expression, a monocyte-specific marker, was lost in cells cultured in the presence of GM-CSF+IL-4 either treated with ESAT-6 or left untreated (Supplementary Fig. S1). Together, these results indicate that while GM-CSF+IL-4 drives the differentiation of CD14 CD209 MoDCs from monocytes, the presence of ESAT-6 impairs this process. ESAT-6–treated cells retained a monocyte-like morphology and failed to express the dendritic cell marker CD209, despite the downregulation of CD14, suggesting a block in differentiation at an intermediate state.

**Figure 1.**
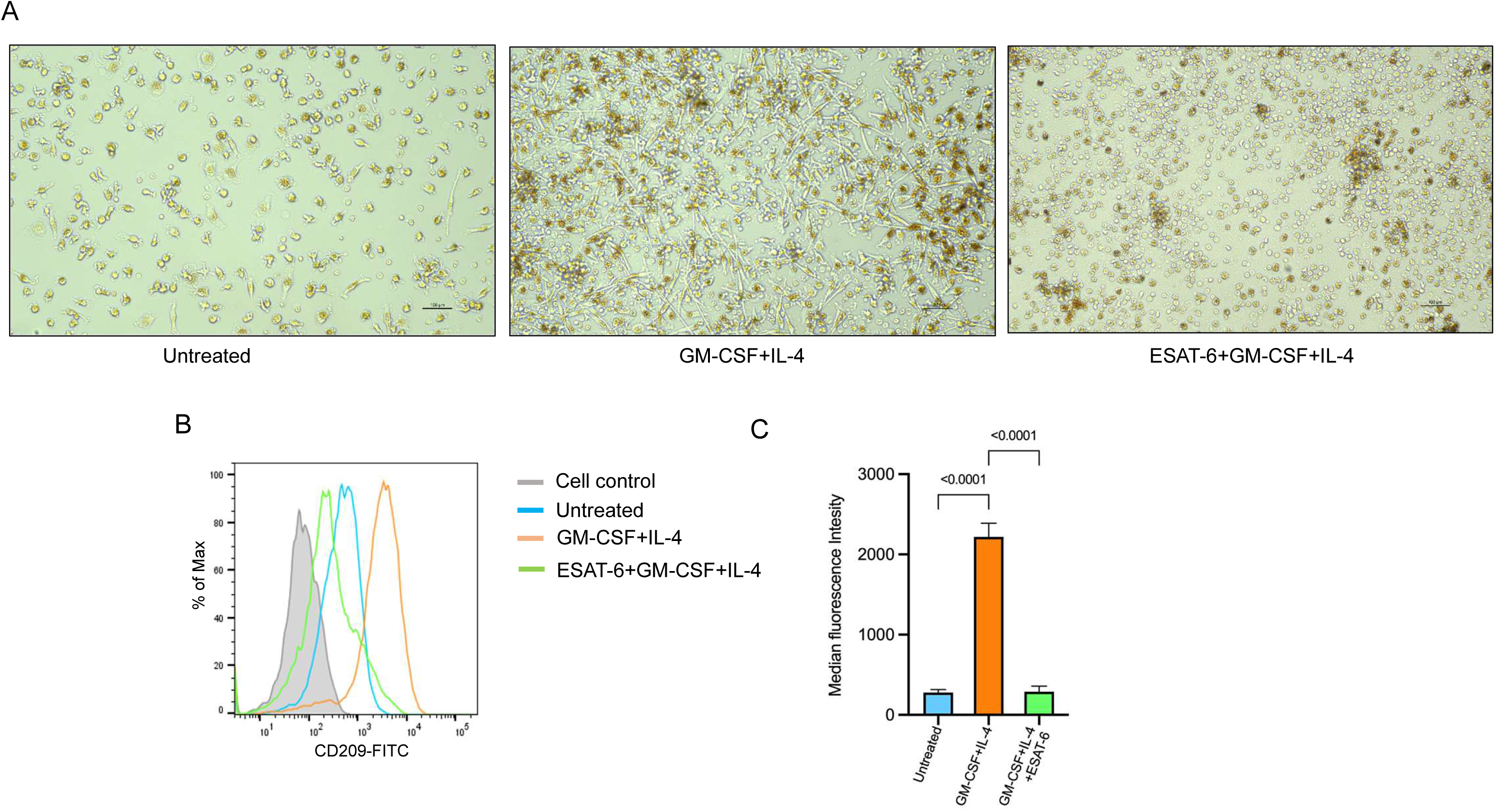
ESAT-6 impairs morphological and phenotypic differentiation of monocytes into dendritic cells. (A) (A) Human peripheral blood monocytes were pre-treated with recombinant ESAT-6 protein (10 µg/ml) for 30 minutes prior to stimulation with GM-CSF (100 ng/ml) and IL-4 (50 ng/ml) to induce differentiation into monocyte-derived dendritic cells (MoDCs). After 7 days, cellular morphology was examined under a phase-contrast microscope. Data are representative of eight independent experiments. Scale bar, 100 µm. (B) Flow cytometry analysis of CD209 (DC-SIGN) expression was performed on day 7. (C) Median fluorescence intensities of different experimental groups were calculated and the results are presented as mean ± SEM of median fluorescence intensity of eight individual samples.

### ESAT-6 induces the expression of IL-6 and IL-10 cytokines in monocyte culture

The cytokines are key determinants in guiding differentiation of monocytes to dendritic cells. Previous studies have demonstrated that cytokines such as IL-6 and IL-10 can direct this process [21,22]. Furthermore, elevated levels of these cytokines have been reported in monocyte cultures infected with Mtb during differentiation into DCs [23]. To investigate whether ESAT-6 regulates cytokine production during monocyte to DC differentiation, we measured IL-6 and IL-10 levels in GM-CSF+IL-4-treated monocyte cultures either left untreated or treated with ESAT-6. As shown in Figure 2, treatment with ESAT-6 resulted in a significant increase in both IL-6 and IL-10 levels compared to untreated controls at both time points. These findings suggest that ESAT-6 induces a cytokine profile characterized by elevated IL-6 and IL-10, which are known to inhibit DC differentiation. Thus, it is possible that ESAT-6 impairs DC differentiation by promoting production of IL-6/IL-10 cytokine.

**Figure 2.**
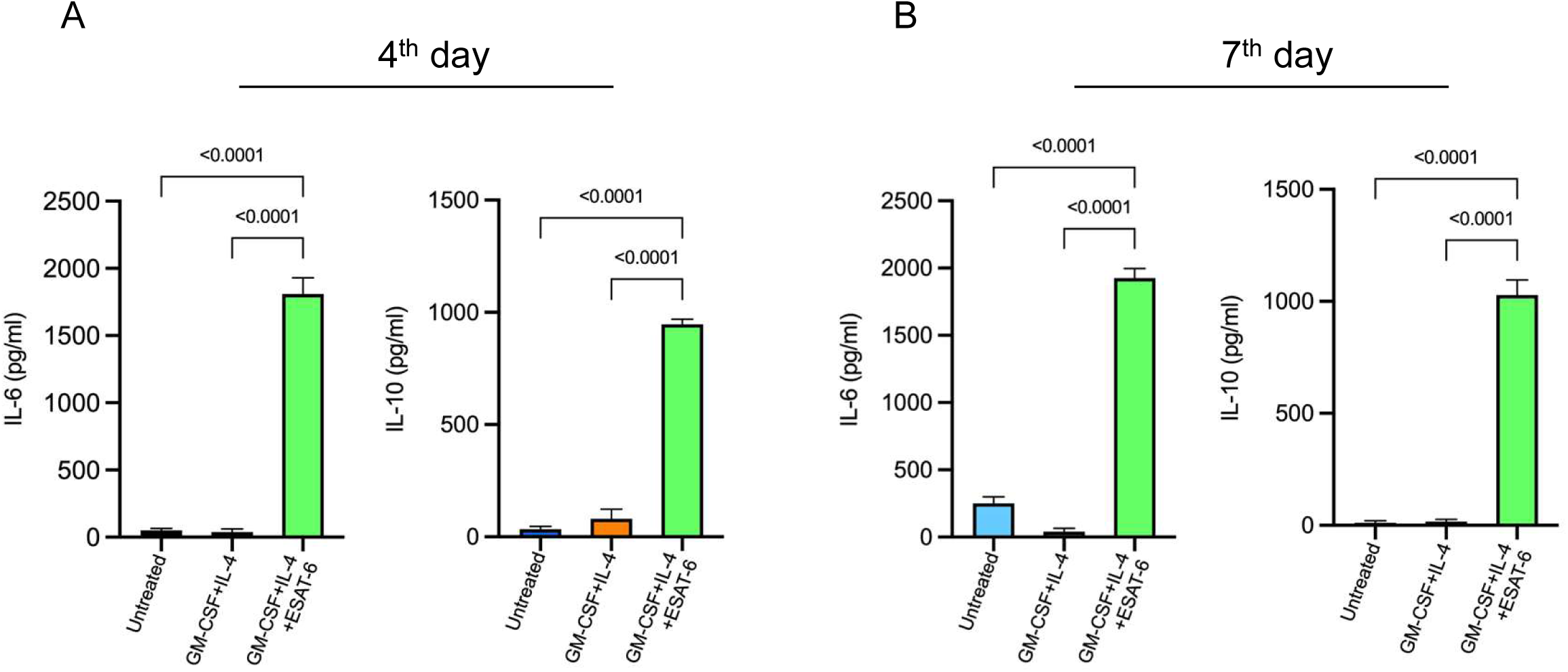
ESAT-6 enhances IL-6 and IL-10 cytokine production during monocyte to dendritic cell differentiation. (A, B) Human monocytes were cultured with GM-CSF+IL-4 to induce differentiation into dendritic cells in the presence or absence of recombinant ESAT-6 (10 µg/ml). Culture supernatants were collected on day 4 (A) and day 7 (B), and levels of IL-6 and IL-10 were quantified by ELISA. Data are presented as mean ± SEM of five individuals.

### IL-6 and IL-10 cytokines induced by ESAT-6 are not responsible for inhibition of DC differentiation from monocytes

Previous studies have demonstrated that both IL-6 and IL-10 are capable of inhibiting the differentiation of DCs from monocytes [21,22]. To investigate whether inhibition of DC differentiation by ESAT-6 is mediated through IL-6/IL-10, we employed specific pharmacological inhibitors and neutralizing antibodies targeting IL-6 and IL-10. GM-CSF+IL-4- treated monocytes were cultured with ESAT-6 in the absence or presence of SB203580, a p38 MAPK inhibitor known to suppress IL-10 production [24] or S3I-201, a STAT-3 inhibitor that blocks IL-10 production [25] as well as IL-6 secretion [26]. Culture supernatants were collected at day 4 to measure IL-6 and IL-10 cytokines by ELISA and cells were harvested at day 7 for checking DC marker (CD209) by flow cytometry. As shown in Fig. 3A and 3B, SB203580 and S3I-201 significantly reduced the levels of IL-6 and IL-10, respectively in the monocyte cultures treated with ESAT-6 without showing any effect in medium control, however, monocyte differentiation into DCs remained impaired (Fig. 3C and 3D; Supplementary Fig. S2). Additionally, we employed neutralizing antibodies against IL-6 and IL-10 to further block their biological activity during monocyte differentiation. Consistent with the pharmacological inhibition data, neutralization of IL-6 and IL-10 also failed to restore DC differentiation in the presence of ESAT-6 (Fig. 3C and 3D). These findings collectively indicate that the inhibitory effect of ESAT-6 on MoDC differentiation is independent of IL-6/IL-10 signaling. Thus, ESAT-6 likely mediates its suppressive effect through alternative signaling pathways.

**Figure 3.**
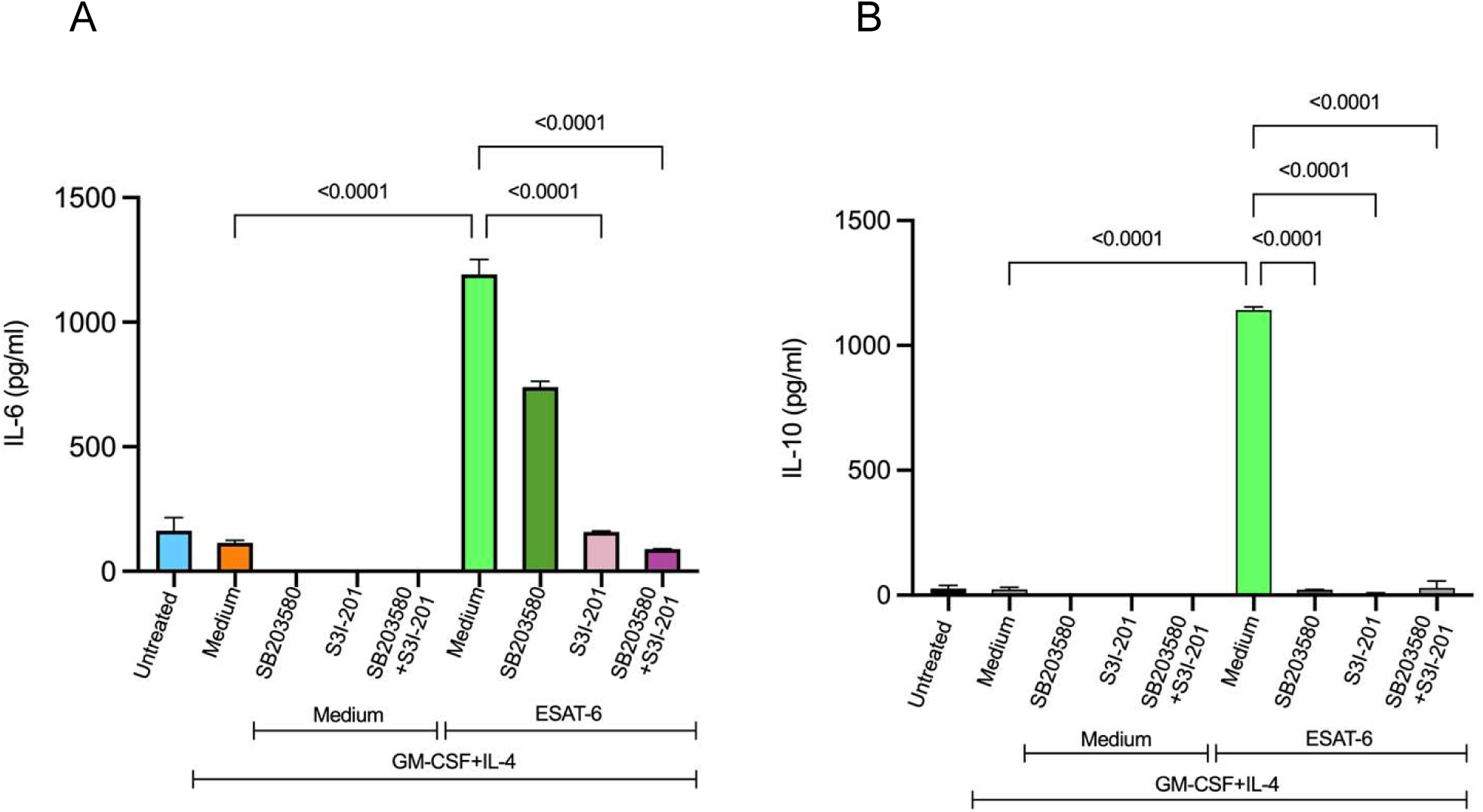
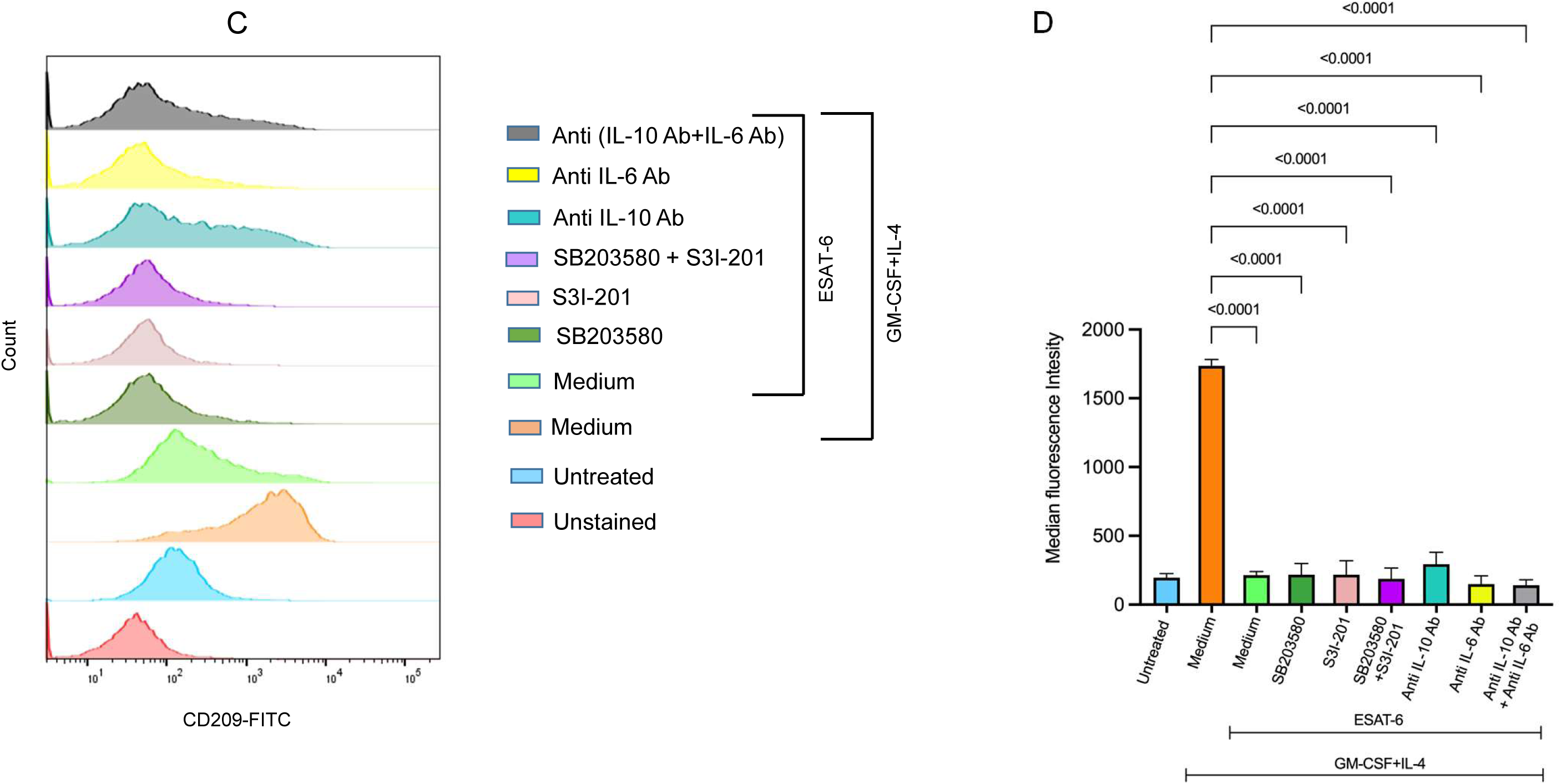
ESAT-6 inhibits dendritic cell differentiation independent of IL-6 and IL-10 signaling. (A) Monocytes were treated with ESAT-6 (10 µg/ml) in the absence or presence of SB203580 (10 µM) or S3I-201 (25 µM), and cultured with GM-CSF+IL-4. Supernatants were collected, and levels of IL-6 (A) and IL-10 (B) were quantified by ELISA. (C) Flow cytometry analysis of DC-specific marker CD209 in monocytes treated with GM-CSF+IL-4 in presence of ESAT-6 or ESAT-6+SB203580 or ESAT-6+S3I-201 or ESAT-6 along with 20 µg/ml of neutralizing antibody against IL-6 or IL-10 during differentiation. (D) Median fluorescence intensities of different experimental groups were calculated. Data represent mean ± SEM of three individual samples.

### ESAT-6 inhibits DC differentiation by inhibiting NF-κB activation

ESAT-6 has previously been reported to inhibit lipopolysaccharide (LPS)-induced activation of NF-κB in macrophages through its interaction with Toll-like receptor 2 (TLR2) and activation of the Akt signaling pathway [13]. Notably, a deletion mutant of ESAT-6 lacking six amino acids at its C-terminus fails to inhibit NF-κB activation. Furthermore, pharmacological inhibition of Akt signaling using wortmannin abrogates the ability of ESAT-6 to suppress NF-κB activation [13]. Given the essential role of NF-κB in dendritic cell differentiation and survival [27,28], we investigated whether ESAT-6 impairs monocyte to DC differentiation through suppression of NF-κB signaling. The transcriptional activity of NF-κB is known to be induced upon phosphorylation of its p65 subunit on serine 276 [29] and NF-κB activation is recorded during MoDC differentiation [30]. Our results show that ESAT-6–treated monocytes had reduced phospho-p65 levels compared to those treated with GM-CSF+IL-4 alone, indicating suppressed NF-κB activation (Fig. 4A) and this was corelated well with impaired DC differentiation (Fig. 4B) and expression of DC marker, CD209 (Fig. 4C and 4D). Importantly, the C-terminal six amino acids deleted mutant ESAT-6 (ΔESAT-6) was unable to suppress NF-κB activity as ΔESAT-6-treated cells exhibited higher phospho-p65 level as compared to ESAT-6-treated cells (Fig. 4A) and the cells showed typical dendritic morphology (Fig. 4B) with increased expression of the DC marker, CD209 (Fig. 4C), which is similar to cells treated with GM-CSF+IL-4 alone. Importantly, treatment with wortmannin (inhibitor of Akt-signaling pathway [13], restored NF-κB activation in the presence of ESAT-6 as these cells exhibited elevated phospho-p65 level, increased surface expression of CD209, and displayed characteristic DC morphology, indicating successful MoDC differentiation (Fig. 4A-D). In contrast, treatment with the NF-κB inhibitor, BAY 11-7082 [31] mimicked the effects of ESAT-6, resulting in decreased phospho-p65 (Fig. 4A), impaired DC morphology (Fig. 4B) and reduced CD209 expression (Fig. 4C-D). These findings collectively demonstrate that ESAT-6 inhibits MoDC differentiation by suppressing p65 NF-κB activation *via* the Akt-dependent pathway. The C-terminal six amino acids of ESAT-6 are found to be critical for this inhibitory effect.

**Figure 4.**
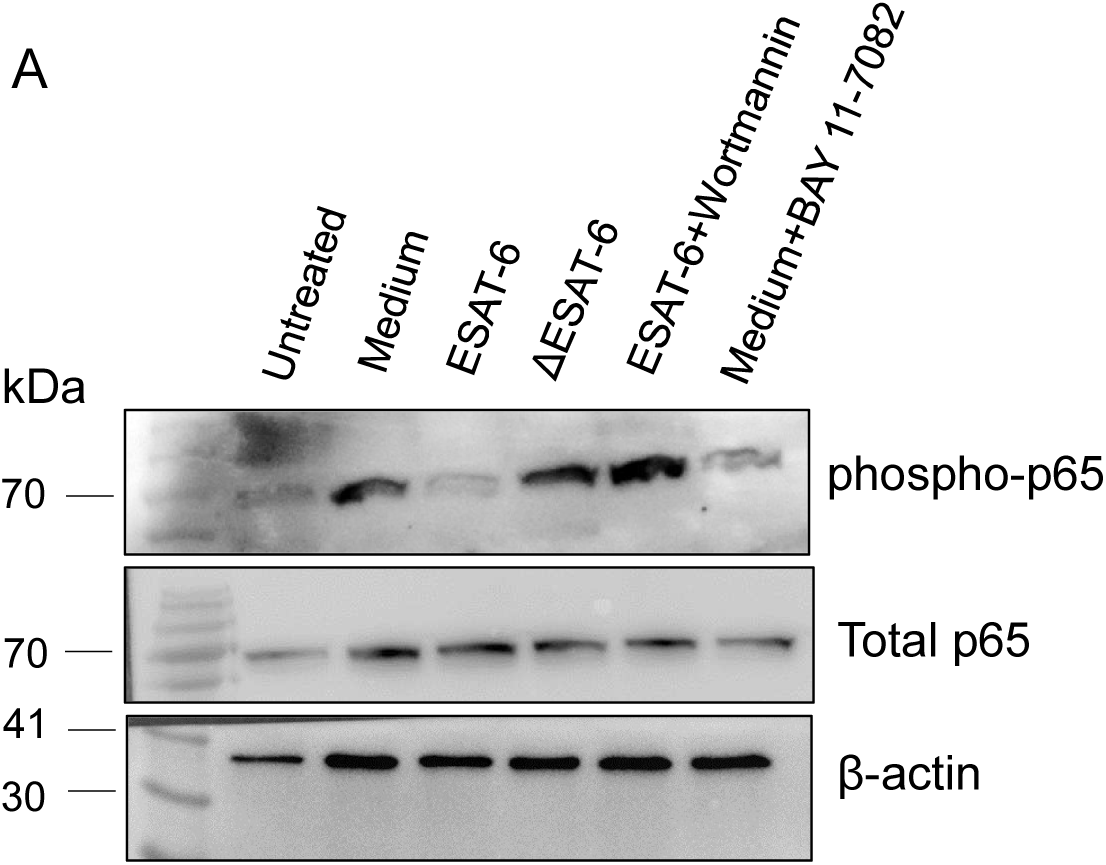
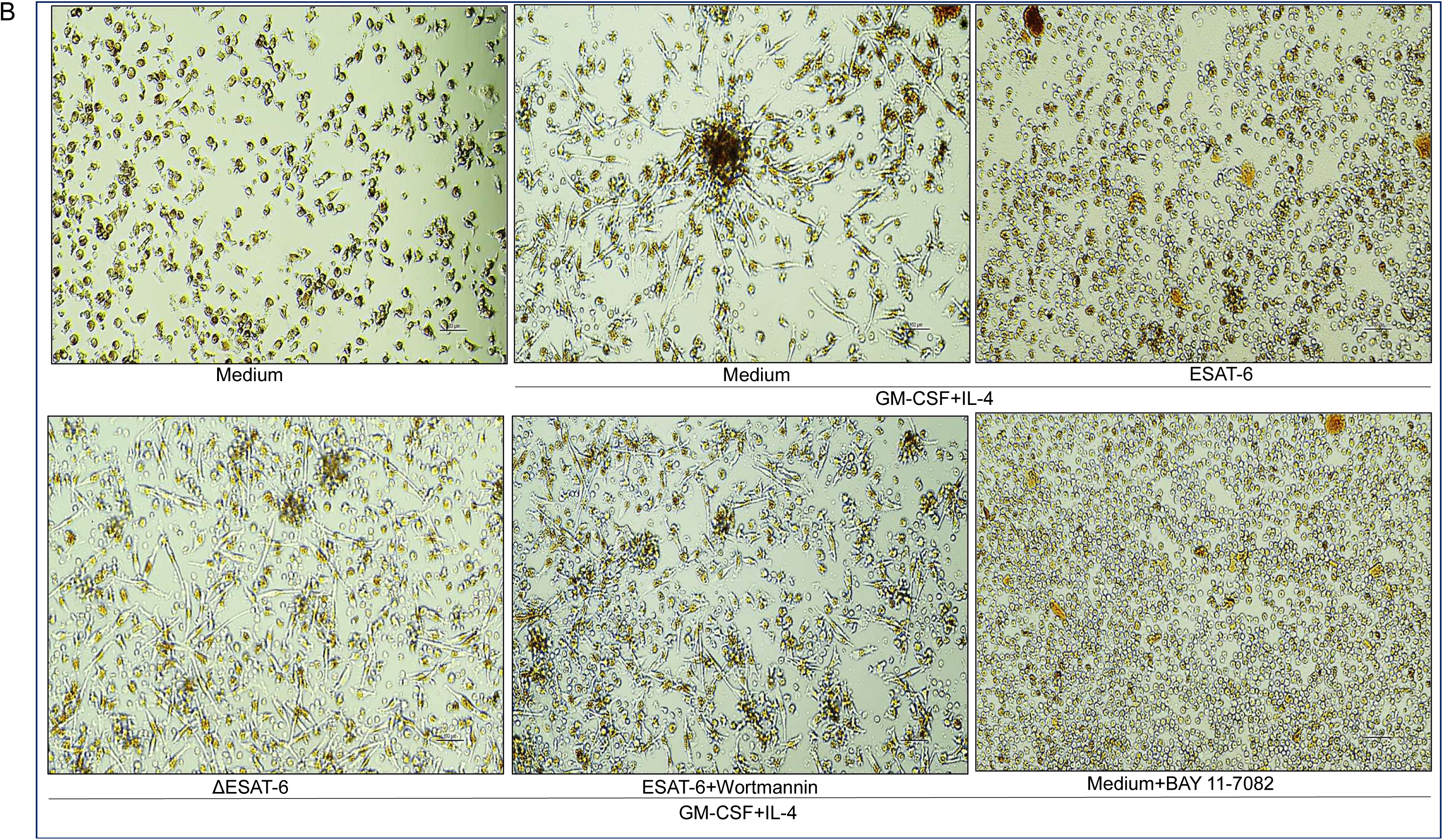
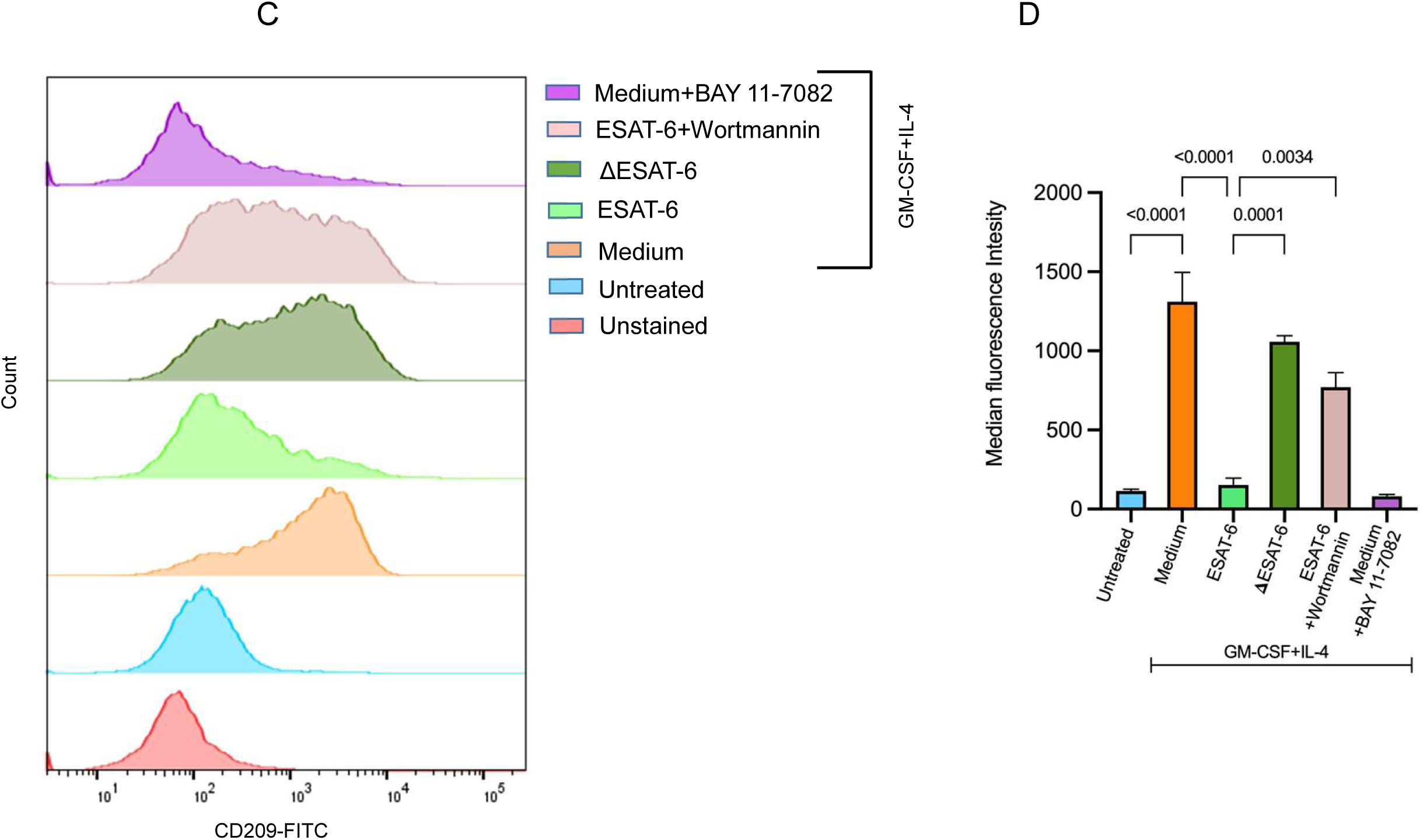
ESAT-6 impairs dendritic cell differentiation by suppressing NF-κB activation. (A) Western blot analysis of phospho-p65 NF-κB was performed on monocyte lysates after 4 days of differentiation with GM-CSF+IL-4 in the absence or presence of either ESAT-6 (10 µg/ml), or ΔESAT-6 (10 µg/ml) or wortmannin (Akt inhibitor, 1 µM), or BAY 11-7082 (NF-κB inhibitor, 10 µM). (B) Cellular morphology was examined under a phase-contrast microscope. Scale bar, 100 μm. (C) Flow cytometric analysis of CD209 in monocyte culture treated with GM-CSF+IL-4 in the absence or presence of ESAT-6, or ΔESAT-6 or wortmannin or BAY 11-7082. Data represent mean ± SEM of three individual samples.

## Discussion

Dendritic cells (DCs) are central to orchestrating host immune responses against Mtb infection [4,32]. They serve as the most potent antigen-presenting cells capable of priming naïve T-cells and initiating anti-Mtb immune responses [4]. DCs are not only derived from bone marrow progenitors but can also be differentiated from circulating monocytes under the influence of specific cytokines such as GM-CSF and IL-4 [7,]. A subset of DCs present in lymph nodes has been shown to originate from peripheral blood monocytes [33], highlighting the plasticity of monocyte differentiation in immune defense. However, Mtb has developed mechanisms to interfere with this critical immune process [33]. We report here that one of the crucial virulent factors that interferes with monocyte-derived DC differentiation is ESAT-6, a known virulence factor that suppresses innate and adaptive immune response [11,13–18].

ESAT-6 was found to interfere with monocyte differentiation into fully competent DC. Monocytes cultured with GM-CSF+IL-4 in the presence of ESAT-6 failed to express the DC-specific marker CD209 (DC-SIGN) [9] and did not develop characteristic dendritic morphology. These findings suggest that ESAT-6 disrupts the differentiation pathway of MoDCs, possibly contributing to the immune evasion strategy of Mtb. The functional consequences of this inhibition are profound, as it may hinder the generation of effective adaptive immunity and the subsequent clearance of the pathogen (11).

Cytokine profiling of ESAT-6-treated cultures revealed increased levels of IL-6 and IL-10, both of which have been implicated in the negative regulation of DC differentiation. Previous studies have shown that IL-10 can inhibit MoDC differentiation and function [21,23], while IL-6 interferes with DC development by skewing monocyte differentiation toward macrophage [22]. Inhibition of p38 MAPK has been shown to downregulate IL-10 production in human monocytes [24] and inhibition of STAT-3 by S3I-201 reduced IL-6 production [24]. However, our results indicate that neither neutralization of IL-6 and IL-10 by specific antibodies nor inhibition of their upstream regulators (STAT3 and p38 MAPK using S3I-201 and SB203580, respectively) was sufficient to rescue the differentiation block, suggesting that additional signaling mechanisms are involved in ESAT-6-mediated immune modulation. One such mechanism involves the transcription factor NF-κB, a critical regulator of DC differentiation [27,28]. Consistent with previous reports showing ESAT-6 can inhibit NF-κB activity *via* activation of Akt signaling [13], we observed reduced p65 phosphorylation in monocytes treated with ESAT-6. Importantly, use of BAY 11-7082, an NF-κB inhibitor [31], recapitulated the differentiation block induced by ESAT-6, supporting the role of NF-κB suppression in this process. Furthermore, we demonstrate that wortmannin, an inhibitor of Akt-signaling pathway [13] was able to reverse the MoDC differentiation inhibition and restore p65 phosphorylation in ESAT-6-treated cultures. This suggests that ESAT-6 acts upstream of NF-κB, potentially through Akt-mediated signaling to suppress DC development. A deletion mutant of ESAT-6 lacking the C-terminal six amino acids (ΔESAT-6) failed to inhibit NF-κB and allowed normal differentiation of MoDCs, reinforcing the importance of this region in mediating immune suppression.

The survival and differentiation of MoDCs are critically dependent on canonical NF-κB signaling, particularly the phosphorylation and nuclear translocation of p65 is essential for the generation of functional MoDCs [27,30]. This pathway is essential not only for DC maturation but also for functional antigen presentation [28]. Consistent with this, inhibition of NF-κB has been shown to impair DC-SIGN (CD209) expression on MoDCs, while downregulation of the monocyte marker CD14 proceeds independently of NF-κB activity [27]. These studies support our observation that ESAT-6–mediated suppression of NF-κB correlates with a block in DC differentiation despite CD14 loss. In summary, these findings highlight a previously uncharacterized function of ESAT-6 in impairing DC differentiation and disrupting early stages of the host immune response. The release of ESAT-6 into the lung microenvironment or systemic circulation during Mtb infection may contribute to immunosuppression by blocking the recruitment and differentiation of monocyte-derived DCs, which are critical for antigen presentation and initiation of T-cell responses [4,9]. Collectively, our study identifies a novel mechanism by which Mtb evades host immunity through ESAT-6-mediated inhibition of NF-κB- dependent DC differentiation. This not only advances our understanding of Mtb pathogenesis but also opens avenues for the development of host-directed therapies in tuberculosis aimed at restoring dendritic cell function targeting the ESAT-6/Akt-NF-κB axis. Targeting the C-terminal domain of ESAT-6 presents a promising therapeutic strategy, as this region is critical for its immune-modulatory functions. Therapeutic interventions aimed at blocking or mimicking this domain could potentially restore normal immune function and counteract *M. tuberculosis* immune evasion, offering a novel approach in host-directed therapy against tuberculosis.

## Acknowledgments

We thank Dr. Vishwanath Jha, Dr Ravi and Dr. Komal Dolasia for suggestion and help. The authors gratefully acknowledge the financial support by the Science and Engineering Research Board (SERB), Department of Science and Technology (DST), Govt of India (JCB/2021/000035), Council of Scientific and Industrial Research (CSIR), Govt. of India (37WS(0020)/2023-24/EMR-II/ASPIRE), Department of Biotechnology (DBT), Govt. of India (BT/PR51149/MED/29/1660/2023), Indian Council of Medical Research (ICMR), Govt. of India (2021-10087/GTGE/ADHOC-BMS and IIRPSG-2024-01-01453) and a core grant from CDFD by DBT. AGM was supported by the fellowship from CSIR. RQ is supported by a grant (R.12013/22/2022-HR) from Department of Health Research (DHR), Govt of India.

## Abbreviations used in this paper

PBMC: Peripheral blood mononuclear cell
DC: Dendritic cell
MoDC: Monocyte-derived dendritic cell
ESAT-6: Early secretory antigenic target of 6 kDa
p38 MAPK: p38 mitogen-activated protein kinase
NF-κB: Nuclear factor kappaB
IL: Interleukin

## References

1. Global tuberculosis report 2024. Geneva: World Health Organization; 2024. Licence: CC BY-NC-SA 3.0 IGO,

2. W. Zhai, F. Wu, Y. Zhang, Y. Fu, Z. Liu, The immune escape mechanisms of *Mycobacterium tuberculosis*, Int. J. Mol. Sci. 20 (2019) 340, 10.3390/ijms20020340.

3. A. Baena, S.A. Porcelli, Evasion and subversion of antigen presentation by *Mycobacterium tuberculosis*, tissue antigens 74 (2009) 189–204, 10.1111/j.1399-0039.2009.01301.x.

4. A. Mihret, The role of dendritic cells in *Mycobacterium tuberculosis* infection, Virulence 3 (2012) 654–659, 10.4161/viru.22586.

5. J. Banchereau, F. Briere, C. Caux, J. Davoust, S. Lebecque, Y.-J. Liu, B. Pulendran, K. Palucka, Immunobiology of dendritic cells, Annu. Rev. Immunol. 18 (2000) 767–811, 10.1146/annurev.immunol.18.1.767.

6. J. Liu, X. Zhang, Y. Cheng, X. Cao, Dendritic cell migration in inflammation and immunity, Cell. Mol. Immunol. 18 (2021) 2461–2471, 10.1038/s41423-021-00726-4.

7. F. Sallusto, A. Lanzavecchia, Efficient presentation of soluble antigen by cultured human dendritic cells is maintained by granulocyte/macrophage colony-stimulating factor plus interleukin 4 and downregulated by tumor necrosis factor alpha, J. Exp. Med. 179 (1994) 1109–1118, 10.1084/jem.179.4.1109.

8. B. León, M. López-Bravo, C. Ardavín, Monocyte-derived dendritic cells formed at the infection site control the induction of protective T helper 1 responses against leishmania, Immunity 26 (2007) 519–531. 10.1016/j.immuni.2007.01.017.

9. C. Cheong, I. Matos, J.-H. Choi, D.B. Dandamudi, E. Shrestha, M.P. Longhi, K.L. Jeffrey, R.M. Anthony, C. Kluger, G. Nchinda, H. Koh, A. Rodriguez, J. Idoyaga, M. Pack, K. Velinzon, C.G. Park, R.M. Steinman, Microbial stimulation fully differentiates monocytes to DC-SIGN/CD209+ dendritic cells for immune T cell areas, Cell 143 (2010) 416–429. 10.1016/j.cell.2010.09.039.

10. B. León, C. Ardavín, Monocyte derived dendritic cells in innate and adaptive immunity, Immunol. Cell Biol. 86 (2008) 320–324, 10.1038/icb.2008.14.

11. A.G. Manikoth, M.K. Bisht, S. Ghosh, S. Mukhopadhyay, Countering the effector functions of ESAT 6 protein in *Mycobacterium tuberculosis*: strategies for developing antimycobacterial therapeutics, FEBS J. (2025), 10.1111/febs.70084.

12. K.N. Lewis, R. Liao, K.M. Guinn, M.J. Hickey, S. Smith, M.A. Behr, D.R. Sherman, Deletion of RD1 from *Mycobacterium tuberculosis* mimics Bacille Calmette-Guérin attenuation, J. Infect. Dis. 53(2003) 1677–1693, 10.1086/345862.

13. S.K. Pathak, S. Basu, K.K. Basu, A. Banerjee, S. Pathak, A. Bhattacharyya, T. Kaisho, M. Kundu, J. Basu, Direct extracellular interaction between the early secreted antigen ESAT-6 of *Mycobacterium tuberculosis* and TLR2 inhibits TLR signaling in macrophages, Nat. Immunol. 8 (2007) 610–618, 10.1038/ni1468.

14. A. Refai, S. Gritli, M.-R. Barbouche, M. Essafi, *Mycobacterium tuberculosis* virulent factor ESAT-6 drives macrophage differentiation toward the pro-inflammatory M1 phenotype and subsequently switches it to the anti-inflammatory M2 phenotype, Front. Cell. Infect. Microbiol. 8 (2018), 10.3389/fcimb.2018.00327.

15. T. Tan, W.L. Lee, D.C. Alexander, S. Grinstein, J. Liu, The ESAT-6/CFP-10 secretion system of *Mycobacterium marinum* modulates phagosome maturation, Cell. Microbiol. 8 (2006) 1417–1429, 10.1111/j.1462-5822.2006.00721.x.

16. H. Dong, W. Jing, Z. Runpeng, X. Xuewei, M. Min, C. Ru, X. Yingru, N. Shengfa, Z. Rongbo, ESAT6 inhibits autophagy flux and promotes BCG proliferation through MTOR, Biochem. Biophys. Res. Commun. 477 (2016) 195–201, 10.1016/j.bbrc.2016.06.042.

17. S. Yang, F. Li, S. Jia, K. Zhang, W. Jiang, Y. Shang, K. Chang, S. Deng, M. Chen, Early secreted antigen ESAT-6 of Mycobacterium tuberculosis promotes apoptosis of macrophages via targeting the MicroRNA155-SOCS1 interaction, Cell. Physiol. Biochem. 35 (2015) 1276–1288, 10.1159/000373950.

18. G. Sreejit, A. Ahmed, N. Parveen, V. Jha, V.L. Valluri, S. Ghosh, S. Mukhopadhyay, The ESAT-6 protein of *Mycobacterium tuberculosis* interacts with beta-2-microglobulin (β2M) affecting antigen presentation function of macrophage, PLoS Pathog. (2014) 10:e1004446. 10.1371/journal.ppat.1004446.

19. A. Udgata, R. Qureshi, S. Mukhopadhyay, Transduction of functionally contrasting signals by two mycobacterial PPE proteins downstream of TLR2 receptors, J. Immunol. 197 (2016) 1776–1787, 10.4049/jimmunol.1501816.

20. N. Romani, D. Reider, M. Heuer, S. Ebner, E. Kämpgen, B. Eibl, D. Niederwieser, G. Schuler, Generation of mature dendritic cells from human blood An improved method with special regard to clinical applicability, J. Immunol. Methods 196 (1996) 137–151, 10.1016/0022-1759(96)00078-6.

21. P. Allavena, L. Piemonti, D. Longoni, S. Bernasconi, A. Stoppacciaro, L. Ruco, A. Mantovani, IL-10 prevents the differentiation of monocytes to dendritic cells but promotes their maturation to macrophages, Eur. J. Immunol.. 28 (2020), 10.1002/(SICI)1521-4141(199801)28:01<359::AID-IMMU359>3.0.CO;2-4

22. P. Chomarat, J. Banchereau, J. Davoust, A.K. Palucka, IL-6 switches the differentiation of monocytes from dendritic cells to macrophages, Nat. Immunol. 1 (2000) 510–514, 10.1038/82763.

23. M.E. Remoli, E. Giacomini, E. Petruccioli, V. Gafa, M. Severa, M.C. Gagliardi, E. Iona, R. Pine, R. Nisini, E.M. Coccia, Bystander inhibition of dendritic cell differentiation by *Mycobacterium tuberculosis* induced IL 10, Immunol. Cell Biol. 89 (2010) 437–446, 10.1038/icb.2010.106.

24. Z.G. Dobreva, L.D. Miteva, S.A. Stanilova, The inhibition of JNK and p38 MAPKs downregulates IL-10 and differentially affects c-Jun gene expression in human monocytes, Immunopharmacol. Immunotoxicol. 31 (2009) 195–201, 10.1080/08923970802626276.

25. J. Sakai, J. Yang, C.-K. Chou, W.W. Wu, M. Akkoyunlu, B cell receptor-induced IL-10 production from neonatal mouse CD19+CD43- cells depends on STAT5-mediated IL-6 secretion, eLife 12 (2023), 10.7554/elife.83561.

26. B.-G. Jung, X. Wang, N. Yi, J. Ma, J. Turner, B. Samten, Early secreted antigenic target of 6-kDa of *Mycobacterium tuberculosis* stimulates IL-6 production by macrophages through activation of STAT3, Sci. Rep. 7 (2017), 10.1038/srep40984.

27. L. Van De Laar, A. Van Den Bosch, S.W. Van Der Kooij, H.L.A. Janssen, P.J. Coffer, C. Van Kooten, A.M. Woltman, A nonredundant role for canonical NF-κB in human myeloid dendritic cell development and function, J. Immunol. 185 (2010) 7252–7261, 10.4049/jimmunol.1000672.

28. A. Hernandez, M. Burger, B.B. Blomberg, W.A. Ross, J.J. Gaynor, I. Lindner, R. Cirocco, J.M. Mathew, M. Carreno, Y. Jin, K.P. Lee, V. Esquenazi, J. Miller, Inhibition of NF-κB during human dendritic cell differentiation generates anergy and regulatory T-cell activity for one but not two human leukocyte antigen DR mismatches, Hum. Immunol. 68 (2007) 715–729, 10.1016/j.humimm.2007.05.010.

29. H. Zhong, R.E. Voll, S. Ghosh, Phosphorylation of NF-κB p65 by PKA stimulates transcriptional activity by promoting a novel bivalent interaction with the coactivator CBP/p300, Mol. Cell 1 (1998) 661–671. 10.1016/s1097-2765(00)80066-0.

30. C. Ammon, K. Mondal, R. Andreesen, S.W. Krause, Differential expression of the transcription factor NF-κB during human mononuclear phagocyte differentiation to macrophages and dendritic cells, Biochem. Biophys. Res. Commun. 268 (2000) 99–105. 10.1006/bbrc.1999.2083.

31. J. Moon, I.J. Moon, H. Hyun, J.M. Yoo, S.H. Bang, Y. Song, S.E. Chang, Bay 11 7082, an NF κB inhibitor, prevents post inflammatory hyperpigmentation through inhibition of inflammation and melanogenesis, Pigment Cell Melanoma Res. (2024), 10.1111/pcmr.13207.

32. K.A. Prendergast, J.R. Kirman, Dendritic cell subsets in mycobacterial infection: Control of bacterial growth and T cell responses, Tuberculosis 93 (2012) 115–122, 10.1016/j.tube.2012.10.008.

33. G.J. Randolph, K. Inaba, D.F. Robbiani, R.M. Steinman, W.A. Muller, Differentiation of phagocytic monocytes into lymph node dendritic cells in vivo, Immunity 11 (1999) 753– 761, 10.1016/s1074-7613(00)80149-1.

