## supplemental Table 1 for "ESAT-6 protein of *Mycobacterium tuberculosis* inhibits differentiation of human monocytes to dendritic cells"

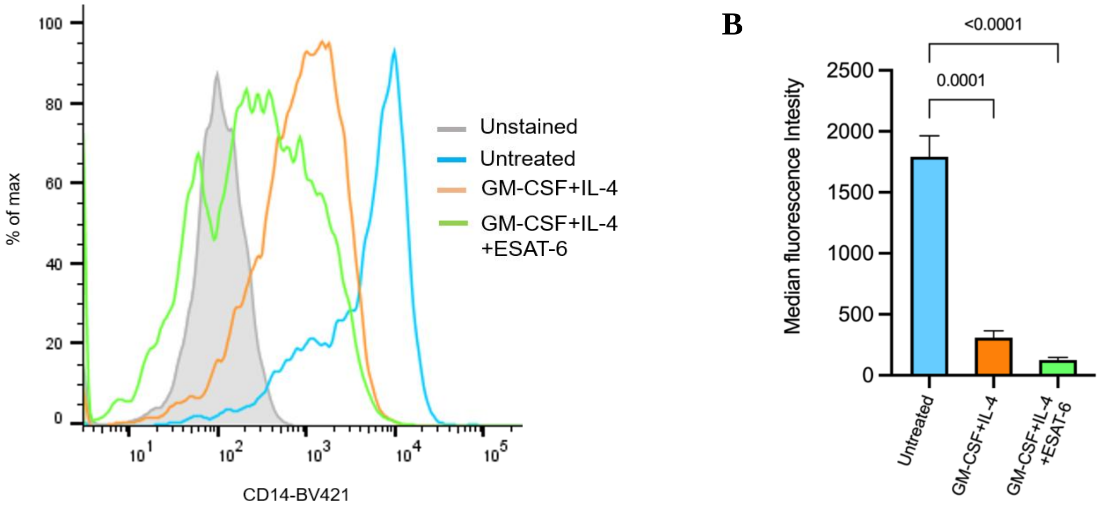


**Supplementary Figure S1: Loss of CD14 marker during differentiation of monocytes to dendritic cells regardless of the presence of ESAT-6.** Human peripheral blood monocytes were pre-treated with recombinant ESAT-6 protein (10 µg/ml) for 30 minutes prior to stimulation with GM-CSF (100 ng/ml) and IL-4 (50 ng/ml) to induce differentiation into monocyte-derived dendritic cells. (A) After 7 days, flow cytometry analysis of CD14 expression was performed using anti-human CD14 Antibody conjugated to Brilliant Violet 421 (CD14-BV421). (B) Median fluorescence intensities of different experimental groups were calculated and the results are presented as mean ± SEM of Median fluorescence intensity of three individual samples. $p < 0.0001$, statistical significance was determined using one-way analysis of variance (ANOVA), followed by Tukey’s multiple comparison test.

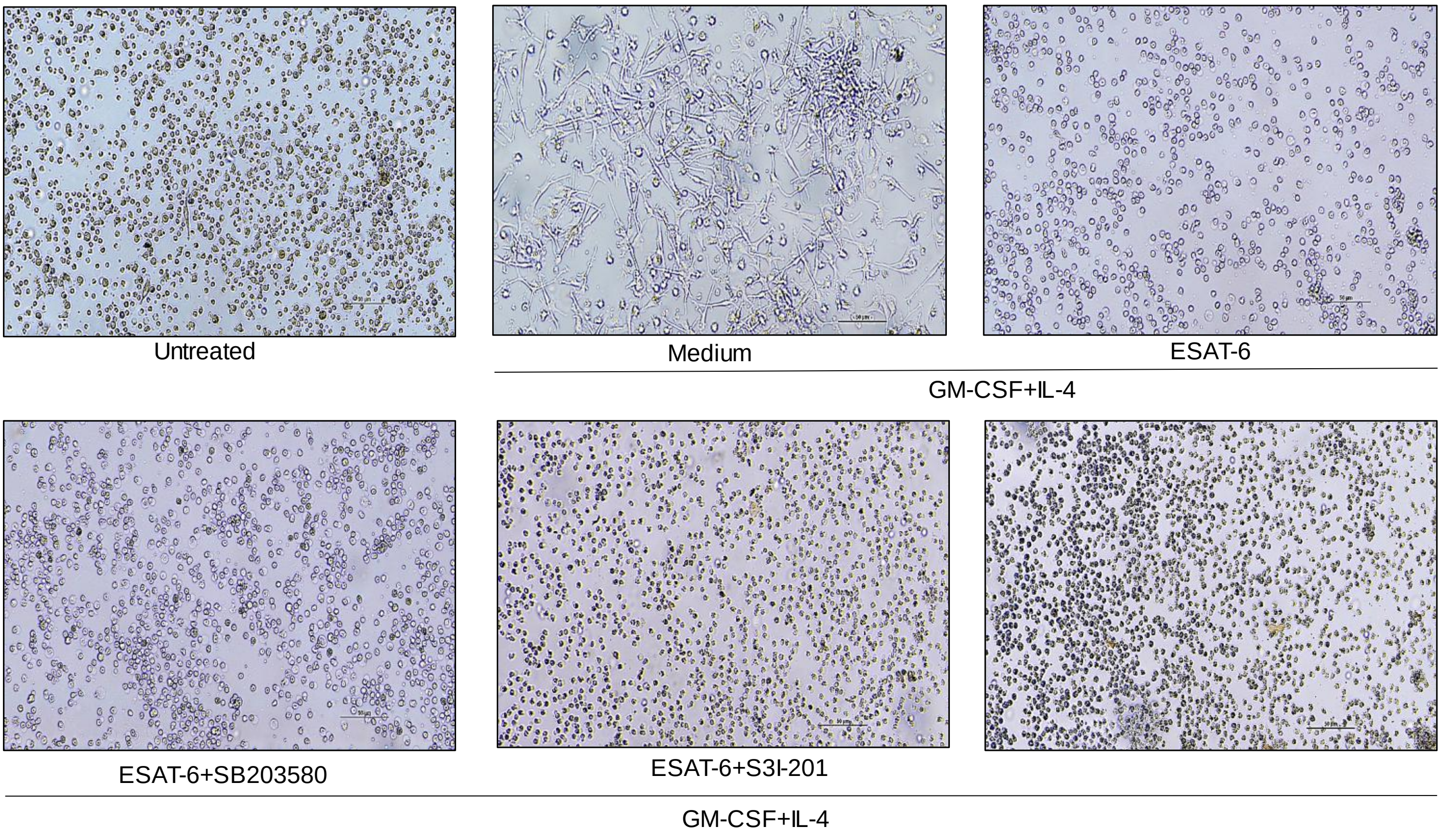


**Supplementary Figure S2: Pharmacological inhibitors of IL-6 and IL-10 fail to rescue ESAT-6 mediated inhibition of DC differentiation.** Human peripheral blood monocytes were pre-treated with recombinant ESAT-6 protein (10 µg/ml) for 30 minutes prior to stimulation with GM-CSF (100 ng/ml) and IL-4 (50 ng/ml) to induce differentiation into monocyte-derived dendritic cells. In some cultures, the cells were treated with either SB203580 (10 µM) or S3I-201 (25 µM) or with SB203580 (10 µM) plus S3I-201 (25 µM). After 7 days, cellular morphology was examined under a phase-contrast microscope. Data are representative of three independent experiments. Scale bar, 50 µm.
